# CLEAN-Contact: Contrastive Learning-enabled Enzyme Functional Annotation Prediction with Structural Inference

**DOI:** 10.1101/2024.05.14.594148

**Authors:** Yuxin Yang, Abby Jerger, Song Feng, Zixu Wang, Christina Brasfield, Margaret S. Cheung, Jeremy Zucker, Qiang Guan

**Author notes:** Contributing authors.

## Abstract

Recent years have witnessed the remarkable progress of deep learning within the realm of scientific disciplines, yielding a wealth of promising outcomes. A prominent challenge within this domain has been the task of predicting enzyme function, a complex problem that has seen the development of numerous computational methods, particularly those rooted in deep learning techniques. However, the majority of these methods have primarily focused on either amino acid sequence data or protein structure data, neglecting the potential synergy of combining of both modalities. To address this gap, we propose a novel **C**ontrastive **L**earning framework for **E**nzyme functional **AN**notation prediction combined with protein amino acid sequences and **Contact** maps (CLEAN-Contact). We rigorously evaluated the performance of our CLEAN-Contact framework against the state-of-the-art enzyme function prediction model using multiple benchmark datasets. Using CLEAN-Contact, we predicted novel enzyme functions within the proteome of *Prochlorococcus marinus* MED4. Our findings convincingly demonstrate the substantial superiority of our CLEAN-Contact framework, marking a significant step forward in enzyme function prediction accuracy.

## 1 Introduction

The crucial role of enzyme function annotation in our understanding of the intricate mechanisms driving biological processes governed by enzymes is widely recognized. The Enzyme Commission (EC) number, a numerical classification system commonly used for enzyme function, is a widely recognized standard in these efforts. The depth of insights provided by the EC number ranges from broad categories of enzyme mechanisms to detailed biochemical reactions through its four hierarchical layers of digits. Traditionally, sequence similarity-based methods, such as the basic local alignment search tool for protein (BLASTP) [1] and HH-suite [2], were largely relied upon for annotating EC numbers. However, in recent times, the deep learning revolution has largely solved the protein structure prediction problem, and it is natural to ask how these twin scientific advances can aid in enzyme function prediction [3].

Predicting enzyme function is not merely an academic classification exercise; it holds immense practical value in systems biology and metabolic engineering, particularly in the construction of genome-scale metabolic models. Such predictive capabilities streamline the process of automating the curation of these models by improving the ability to predict which proteins are responsible for observed growth phenotypes under diverse nutrient conditions and distinct genetic backgrounds. Furthermore, precise knowledge of a genome’s metabolic capabilities enables the design of microbial cell factories to fit for purpose to the metabolic engineering goal, be it medicine, biomanufacturing, or bioremediation.

Currently, the majority of deep learning-based models developed for predicting EC numbers focus on either amino acid sequence or structural data of the enzyme. Studies such as that of [4], for example, rely solely on amino acid sequence data for EC number prediction. Conversely, the work in [5] employs a single convolutional neural network to predict EC numbers, focusing more on the structural data. Additionally, a different approach integrating enzyme structure data into the training process is seen in [6], which provides a more comprehensive view of the enzyme’s functionality. The recent addition of a novel method grounded on contrastive learning has also escalated the performance of EC number prediction [7].

Building upon this groundwork, we propose CLEAN-Contact, a contrastive learning framework that amalgamates both amino acid sequence data and protein structure data for superior enzyme function prediction. Our CLEAN-Contact framework has shown a notable improvement over the current state-of-the-art, CLEAN [7], under a variety of test conditions, further emphasizing the potential for combining protein sequence and structure-based deep learning in enzyme function predictive practices.

## 2 Results

### 2.1 Contrastive Learning Framework for Enzyme Functional Annotation Prediction Combined with Protein Contact Maps

We develop a deep learning framework aimed at predicting EC numbers. This framework integrates a protein language model (ESM-2) [8] and a computer vision model (ResNet50) [9] (See Section 4). Protein language models excel at processing and extracting pertinent information from protein amino acid sequences, while computer vision models, especially convolutional neural networks (CNNs), demonstrate superior efficacy in handling image-like data, making CNNs well-suited for extracting relevant information from the square matrix structure of protein contact maps. Among protein language models, ESM-2 stands out with several advantages over its peers like Prot-Bert [10] and ESM-1b [11]. These include more advanced model architectures, larger training dataset, and superior benchmark performance on the 14th Critical Assessment of protein Structure Prediction (CASP14) [12] and Continuous Automated Model Evaluation (CAMEO) [13]. For the computer vision component, ResNet-50 offers an optimal balance between computational efficiency and performance in relevant tasks.

ESM-2 operates as a feedforward neural network, extracting function-aware sequence representations from input protein amino acid sequences. Meanwhile, ResNet50 functions as a feedforward neural network, extracting structure representations from input 2D contact maps derived from protein structures. The CLEAN-Contact framework plays a key role by combining these sequence and structure representations and employing contrastive learning to learn the prediction of EC numbers. The CLEAN-Contact framework consists of three key components: (1) The representations extraction segment (Fig. 1a), which is designed to extract structure representations from contact maps using ResNet50 [9] and sequence representations from amino acid sequences using ESM2 [8]. (2) The contrastive learning segment (Fig. 1b), where contrastive learning is performed to minimize the embedding distances between enzymes sharing the same EC number while maximizing the embedding distances between enzymes with different EC numbers. Specifically, structure and sequence representations are transformed to the same embedding space, leading to the same dimension for both structure and sequence representations. Subsequently, the combined representations are produced by adding the structure and sequence representations in the same embedding space together. The combined representations are used to measure embedding distances between enzymes. And (3) the EC number prediction segment (Fig. 1c), which is responsible for determining the EC number of a query enzyme based on the projector and the combined representation of the query enzyme learned through contrastive learning.

**Fig. 1.**
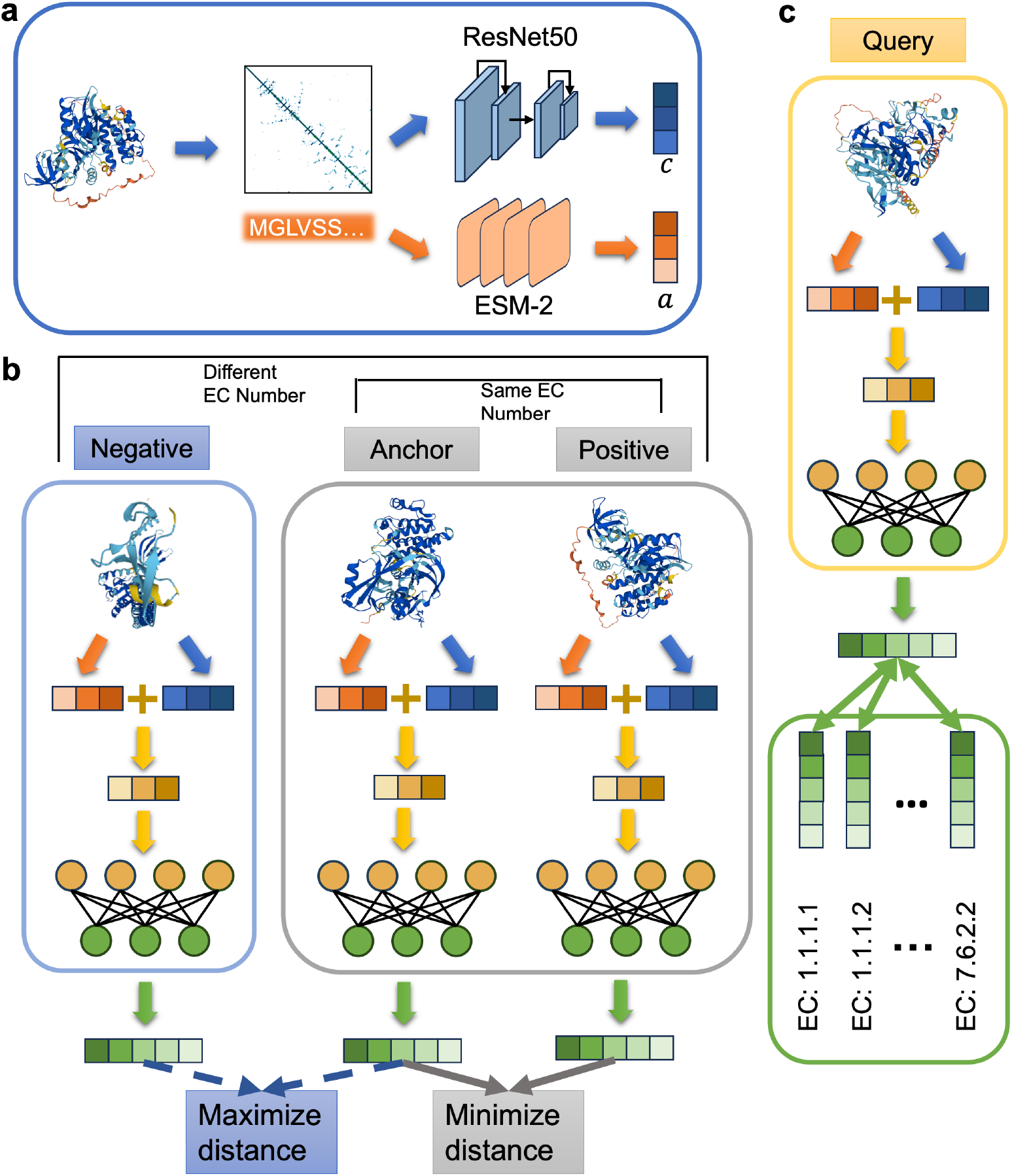
Schematic illustration of the CLEAN-Contact framework. **a**, Obtaining structure and sequence representations from contact map and amino acid sequence using ResNet50 and ESM2, respectively. **b**, The contrastive learning segment. Sequence and structure representations are combined and projected into high-dimensional vectors using the projector. Positive samples are those with the same EC number as the anchor sample and negative samples are chosen from EC numbers with cluster centers close to the anchor. We perform contrastive learning to minimize distances between anchor and positive samples, and maximize distances between anchor and negative samples. **c**, The EC number prediction segment. Cluster centers are computed for each EC number by averaging learned vectors within that EC number. Euclidean distances between the query enzyme’s vector and the cluster centers are calculated to predict the EC number of a query enzyme. *P* -value EC number selection algorithm is used to determine predicted EC numbers.

### 2.2 Benchmark results

We conducted comprehensive evaluations of our proposed CLEAN-Contact framework, comparing it against five state-of-the-art EC number prediction models, CLEAN [7] DeepECtransformer [14], DeepEC [4], ECPred [15], and ProteInfer [5]. These models, along with the CLEAN-Contact, underwent testing on two independent test datasets. The first test dataset, New-392 [7] contains 392 enzyme sequences distributed over 177 different EC numbers. Predictive performance was assessed using four different metrics, Precision, Recall, F1-score, and Area Under Receiver Operating Characteristic Curve (AUC). The CLEAN-Contact achieved better performance performance. Specifically, CLEAN-Contact exhibited a 16.22% enhancement in Precision (0.652 vs. 0.561), a 9.04% improvement in Recall (0.555 vs. 0.509), a 12.30% increase in F1-score (0.566 vs. 0.504), and a 3.19% elevation in AUC (0.777 vs. 0.753) over CLEAN (Fig. 2**a**). Conversely, ECPred and DeepEC recorded the lowest performance.

**Fig. 2.**
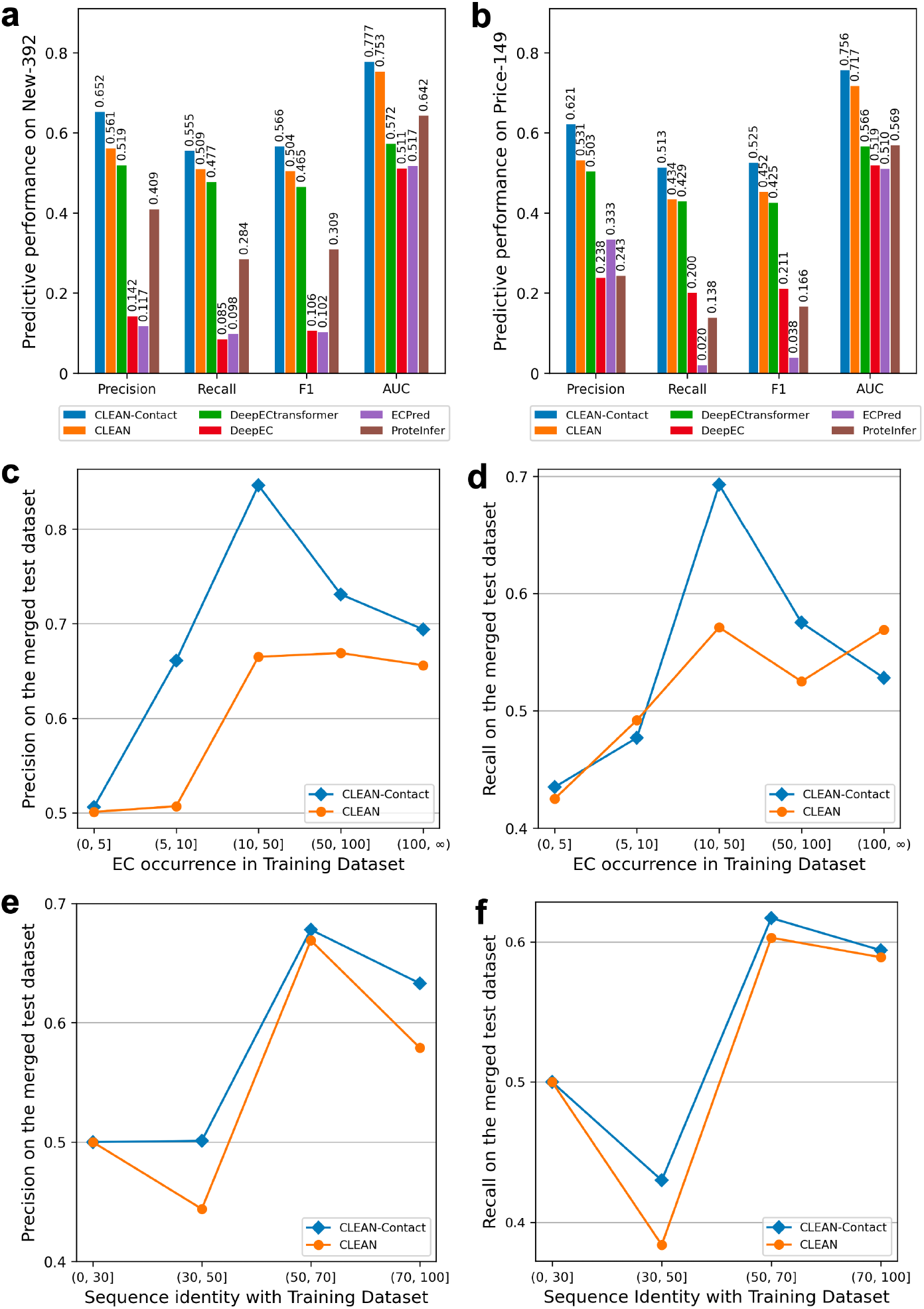
Assessment of predictive performance between CLEAN-Contact and baseline models. **a, b**, Predictive performance between CLEAN-Contact and baseline models (CLEAN, DeepECtransformer, DeepEC, ECPred, and ProteInfer) measured by Precision, Recall, F1-score, and AUC metrics on the New-392 test dataset. **c, d**, Precision and recall of CLEAN-Contact and the second best performing model, CLEAN, on the merged test dataset, correlating with the frequency of occurrence of EC numbers in the training dataset. **e, f**, Precision and recall of CLEAN-Contact and CLEAN on the merged dataset, correlating with the maximum sequence identity of proteins in the test dataset compared to the training dataset.

The second test dataset, Price-149 [7] comprises 149 enzyme sequences distributed over 56 different EC numbers. Once more, CLEAN-Contact exhibited superior performance. Specifically, CLEAN-Contact showcased enhancements across various metrics compared to CLEAN, the second best performing model: a 16.95% improvement in Precision (0.621 vs. 0.531), an 18.20% increase in Recall (0.513 vs. 0.434), a 16.15% boost in F1-score (0.525 vs. 0.452), and a 5.44% elevation in AUC (0.756 vs. 0.717; Fig. 2**b**). ECPred recorded the lowest performance measured by Recall and F1-score, while DeepEC recorded the lowest performance measured by Precision. Notably, CLEAN-Contact achieved a 2.0-to 2.5-fold improvement in Precision compared to DeepEC (0.238), ECPred (0.333), and ProteInfer (0.243), a 25.6-fold increase in Recall compared to ECPred (0.020), and a 13.8-fold improvement in F1-score compared to ECPred (0.038).

To investigate the performance of CLEAN-Contact concerning understudied EC numbers, we merged the two test datasets and divided the merged test dataset based on the frequency of an EC number’s occurrence in the training dataset. Subsequently, we measured the Precision and Recall of both CLEAN-Contact and CLEAN on these divided test datasets. CLEAN-Contact demonstrated a 30.4% improvement (0.661 vs. 0.507) in Precision while achieving comparable performance against CLEAN when the EC number was rare in the training dataset (occurring more than 5 times but less than 11 times) and a 27.4% improvement (0.847 vs. 0.665) in Precision and a 21.4% improvement (0.693 vs. 0.571) in Recall compared to CLEAN when moderately infrequent (occurring more than 10 times but less than 51 times; Fig. 2**c, d**). However, when the EC number was extremely rare in the training dataset (occurring less than 6 times) or very common (occurring more than 100 times), the improvement of CLEAN-Contact over CLEAN was less significant (0.506 vs. 0.501 in Precision and 0.435 vs. 0.425 in Recall for EC numbers occurring less than 6 times, 0.731 vs. 0.669 in Precision and 0.575 vs. 0.525 in Recall for EC numbers occurring more than 50 times but less than 101 times, and 0.694 vs. 0.656 in Precision and 0.528 vs. 0.569 in Recall for EC numbers occurring more than 100 times; Fig. 2**c, d**). These findings underscore the significant enhancement in predictive performance achieved by integrating structural information into CLEAN-Contact.

Furthermore, we divided the merged test dataset based on the maximum sequence identity with the training dataset and evaluated the Precision and Recall of both CLEAN-Contact and CLEAN on these divided test datasets. When the sequence identity with the training dataset was very low (less than 30%), both models achieved comparable performance (Fig. 2**e, f**). However, as the maximum sequence identity ranged from 30% to 50%, CLEAN-Contact exhibited a 12.3% improvement in Precision (0.501 vs 0.446) and a 12.0% improvement in Recall (0.430 vs. 0.384) over CLEAN (Fig. 2**e, f**). As the maximum sequence identity increased to between 50% and 70%, CLEAN-Contact still maintained a 1.34% advantage in Precision (0.678 vs. 0.669) and a 2.32% advantage in Recall (0.617 vs. 0.603) over CLEAN (Fig. 2**e, f**). Remarkably, when the maximum sequence identity exceeded 70%, CLEAN-Contact showcased a 9.33% improvement over CLEAN (0.633 vs. 0.579) in Precision and a comparable performance in Recall (0.594 vs. 0.589; Fig. 2**e, f**).

### 2.3 Discovery of unknown functions of enzymes in *Prochlorococcus marinus* MED4

We next aimed to uncover unknown enzyme functions within the proteome of *Prochlorococcus marinus* (*P. marinus*) MED4 (UniProt Proteome ID: UP000001026). *P. marinus* is a dominant photosynthetic organism in tropical and temperate open ocean ecosystems and is notable for being the smallest known photosynthetic organism [16]. The proteome of *P. marinus* MED4 comprises 1,942 proteins, of which 583 had been annotated with at least one EC number in the UniProt database [17], which encompasses a total of 488 distinct EC numbers.

We first employed both CLEAN-Contact and CLEAN to predict EC numbers for all 1,942 proteins in the *P. marinus* MED4 proteome. Of the 488 annotated EC numbers, CLEAN-Contact correctly predicted 385, while CLEAN correctly predicted 379 (Fig. 3**a** and Supplementary Data 1). Both methods predicted 373 new EC numbers, with CLEAN-Contact independently predicting an additional 442 new EC numbers (Fig. 3**a**). CLEAN-Contact had higher overall prediction confidence compared to CLEAN (cf. Section S1.5), further confirming the better performance of CLEAN-Contact through integrating both protein structure and sequence information (Fig. 3**b**). Analysis of the first-level EC numbers predicted by CLEAN-Contact revealed that EC:2 comprised the most of predictions (30.2%), followed by EC:1 (21.5%) and EC:3 (21.3%), with the remaining first-level EC numbers accounting for 27% (Fig. 3**c**).

**Fig. 3.**
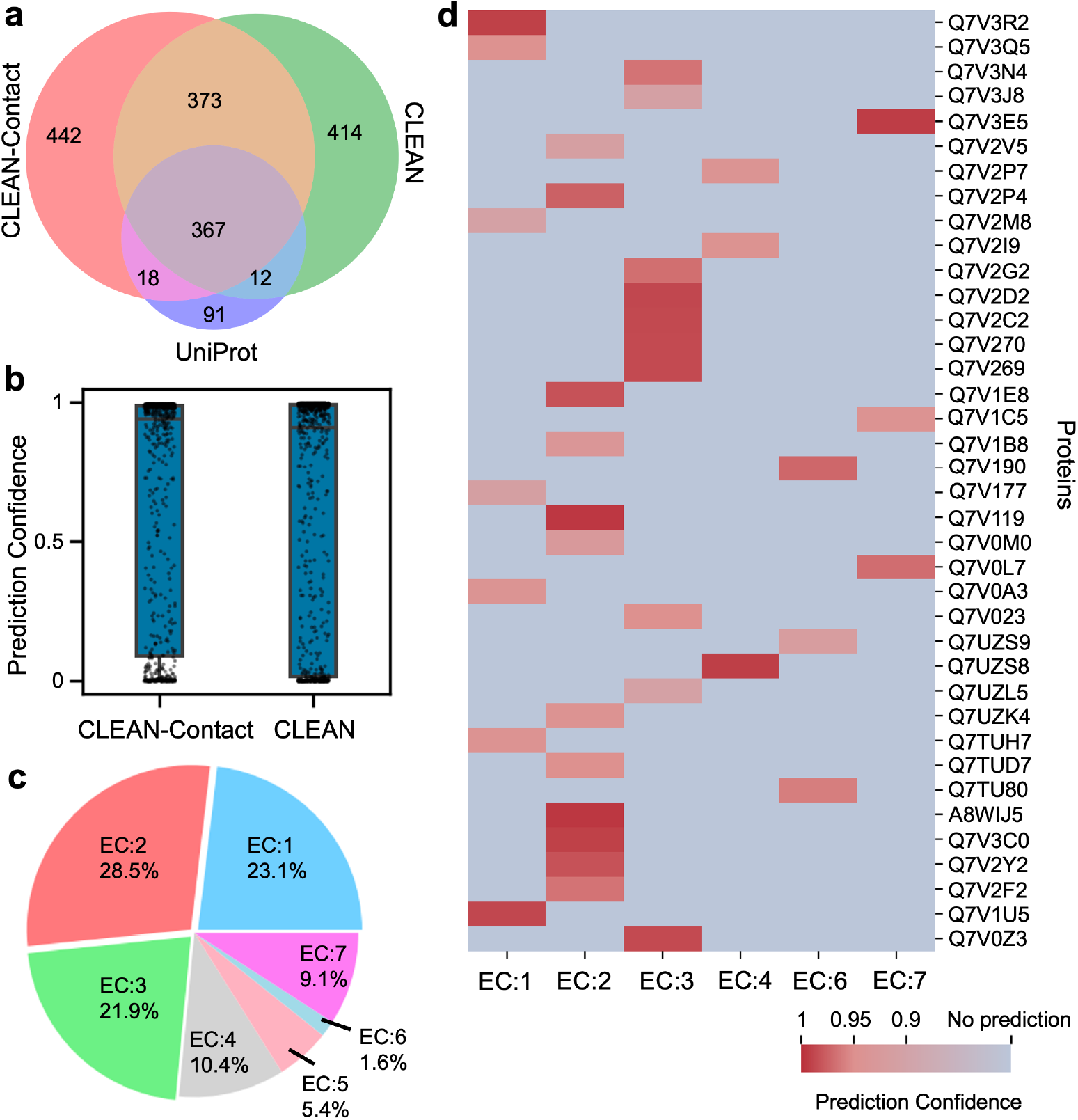
Enzyme function prediction within the proteome of *Prochlorococcus marinus* MED4. **a**, Evaluation of predictive performance between CLEAN-Contact and CLEAN on 583 annotated enzymes. **b**, Assessment of prediction confidence between CLEAN-Contact and CLEAN for the 583 annotated enzymes. Box plot elements: The center lines represent the median prediction confidences. The upper and lower box limits represent upper and lower quartiles, respectively. Whiskers extend to the maximum and minimum prediction confidences. Individual points represent all prediction confidence values. **c**, Distribution of first-level EC numbers predicted by CLEAN-Contact for 1,212 unannotated and unreviewed enzymes. **d**, Visualization of 38 enzymes with high-confidence predictions and their corresponding predicted first-level EC number.

We next focused our attention on the predicted EC numbers for proteins lacking annotated EC numbers and in the “unreviewed” status, indicating they had not yet been manually annotated by experts. A total of 1,212 proteins meet these criteria. To ensure prediction reliability, we applied strict confidence threshold. By considering only predictions with a confidence score exceeding 0.9, we identified 38 enzymes with a total of 36 predicted EC numbers (Fig. 3**d** and Supplementary Data 2). Notable examples include protein with UniProt ID of Q7V3C0, predicted as citrate synthase (EC:2.3.3.16) with 0.991 confidence, aligning with its protein name in the UniProt database, protein with UniProt ID of A8WIJ5, uncharacterized protein in the UniProt database, predicted as tetrahydromethanopterin S-methyltransferase (EC:2.1.1.86) with 0.996 confidence, and protein with UniProt ID of Q7V190, conserved hypothetical protein in the UniProt database, predicted to be 6-carboxyhexanoate–CoA ligase (EC:6.2.1.14) with 0.972 confidence (Fig. 3**d** and Supplementary Data 2). Together, these results demonstrated CLEAN-Contact’s potential for identifying unknown enzyme functions, particularly uncharacterized or hypothetical proteins.

### 2.4 Interpreting the neural network

We investigated the impact of various representation components, specifically comparing sequence representations derived from ESM-2 against those derived from ESM-1b, along with including structure representations. We observed that replacing ESM-1b with ESM-2, without incorporating structure representations, led to a marginal 1.39% average performance improvement across the two test datasets (Fig. 4). However, integrating structure representations while retaining ESM-1b yielded a substantial 5.76% average performance increase across the two test datasets (Fig. 4). Moreover, replacing ESM-1b with ESM-2 and including structure representations resulted in a 6.13% average performance improvement (Fig. 4). Notably, utilizing solely structure representations as the model input yielded the poorest performance. We attributed this outcome to the fact that contact maps only offer information about residue contacts within the protein structure, lacking crucial details about the amino acids.

**Fig. 4.**
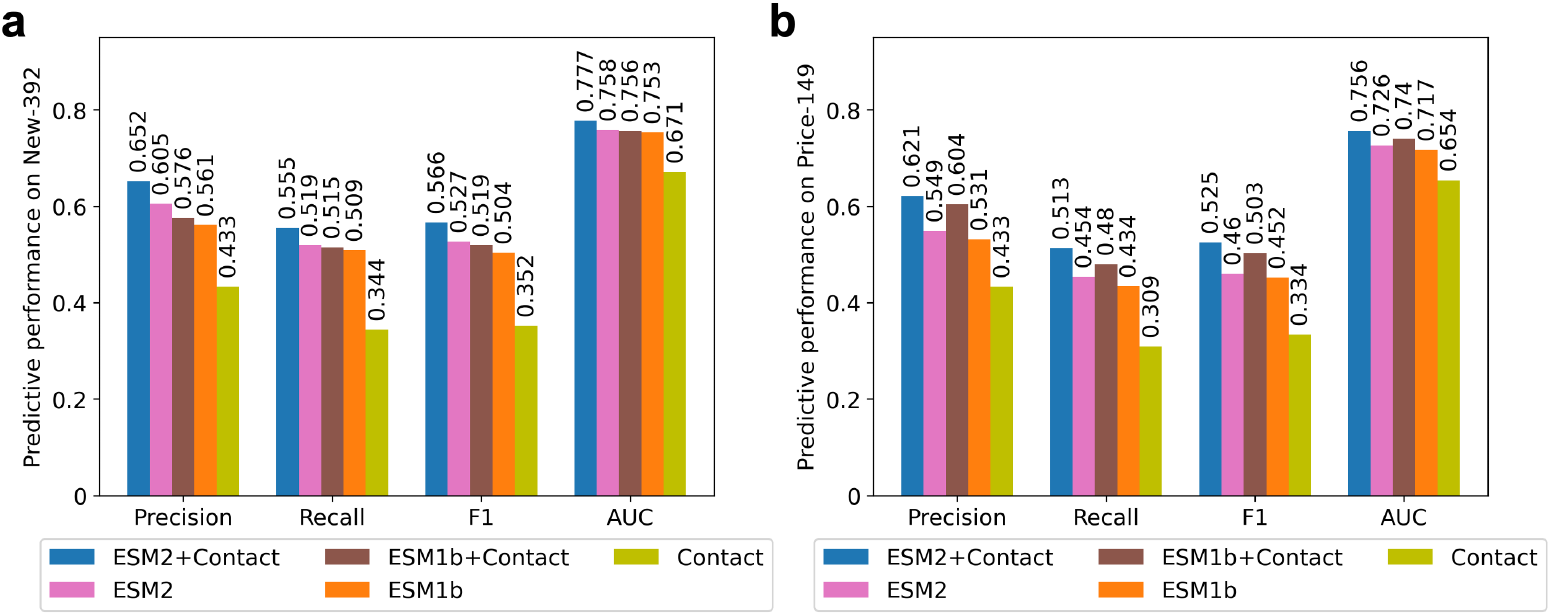
Ablation results. We conducted a comparative analysis of several model variations, including our proposed CLEAN-Contact (ESM2+Contact), CLEAN-Contact without structure representation (ESM2), CLEAN-Contact with ESM-1b instead of ESM-2 (ESM1b+Contact), the original CLEAN (ESM1b), and a model with solely structure representation devoid of ESM-2 or ESM-1b generated sequence representation (Contact). **a**, Evaluation of predictive performance metrics (Precision, Recall, F-1 score, and AUC) across different models on the New-392 test dataset. **b**, Evaluation of predictive performance metrics (Precision, Recall, F-1 score, and AUC) across different models on the Price-149 test dataset.

## 3 Discussion

In this work, we introduce a novel framework based on contrastive learning, which integrates both enzyme amino acid sequence and structural data to predict EC numbers. Our proposed CLEAN-Contact framework harnesses the power of ESM-2, a pretrained protein language model responsible for encoding amino acid sequences, and ResNet, a convolutional neural network utilized for encoding contact maps. Through comprehensive evaluations on diverse test datasets, we have meticulously assessed the CLEAN-Contact framework’s performance. In addition to benchmark analysis, we leveraged CLEAN-Contact to discover novel enzyme functions in *Prochlorococcus marinus* MED4. Specifically, CLEAN-Contact predicted enzyme functions with high confidence for 38 proteins that had not been manually annotated and reviewed by experts. Our extensive comparisons and detailed analyses firmly established that the fusion of structural and sequence information substantially enhances the predictive performance of models used for enzyme functional annotation prediction. As a result, CLEAN-Contact represents a significant step forward in the field of enzyme annotation, providing a robust framework for enzyme function prediction.

However, our work does come with certain limitations. First, our utilization of structure information relies on contact maps a 2D matrix representation, rather than utilizing the full 3D protein structures of enzymes. One potential solution to overcome this limitation involves the incorporation of 3D interaction sequences (3Di) [18]. These sequences contain valuable information regarding geometric conformations between residues. Another possible way to enhance CLEAN-Contact involves considering EC numbers as hierarchical labels. This approach entails employing hierarchical losses, such as hierarchical contrastive loss [19] or exploring other hierarchical classification loss methodologies [20, 21].

Even though our focus in this work is solely on predicting enzyme functional annotations, we firmly believe that our proposed CLEAN-Contact framework has broader applications beyond this domain. A promising future direction could involve extending our model to predict general protein functional annotations, such as Gene Ontology (GO) numbers [22] and FunCat categories [23]. This expansion would significantly broaden the application and utility of our model.

## 4 Methods

### 4.1 Description of Dataset

The enzyme’s amino acid sequences in the training dataset were retrieved from Swiss-Prot [24] released in April 2022. Sequences lacking structures available in the AlphaFold Protein Structure Database [25] (https://alphafold.ebi.ac.uk/) were filtered out from the training dataset. The processed dataset comprises 224,742 amino acid sequences, covering 5,197 EC numbers. Test datasets, New-392 and Price-149, consist of 392 and 149 amino acid sequences, respectively, distributed across 177 and 56 EC numbers, as provided by [7] (See Table S5 and S6). For enzymes in the test datasets without available structures, we generated protein structures using AlphaFold2 [3]. The case study dataset, comprising 1,942 proteins, was obtained from the UniProt database (UniProt proteome ID: UP000001026, accessed September 2024). Of 1,942 proteins, 583 were annotated with at least one EC number, while 1,212 were neither annotated nor reviewed by experts. There is no overlapping between the proteins in the case study dataset and those in the training dataset.

### 4.1 Description of Framework

As shown in Fig. 1, our framework consists of two components: the contrastive learning segment and the EC number prediction segment.

We initiate our approach by obtaining contact maps 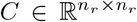 from protein structures, where *n*_*r*_ denotes the number of residues in a protein. To expand *C* into a three-channel matrix 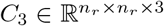, each channel holds identical values. Utilizing the ResNet50 [9] model pretrained on ImageNet [26], we extract high-dimensional structure representations from contact maps:

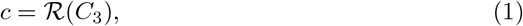

where *ℛ* is the pretrained ResNet50 without its classification layer, and 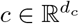 is the structure representation, with *d*_*c*_ as its dimension.

Recognizing that contact maps alone lack vital amino acid information, we leverage ESM-2 [8] to derive sequence representations from proteins’ amino acid sequence:

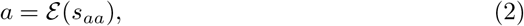

where *ε* is the pretrained ESM-2 model, *s*_*aa*_ is the protein’s amino acid sequence, and 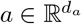 is the sequence representation, with *d*_*a*_ as its dimension.

To fuse the sequence and structure representations into a vector 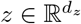, we employ a projector *F*_Ψ_(·). The projector contains three levels of linear layers, each followed by layer normalization [27]:

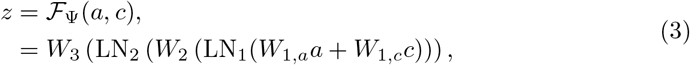

where LN_*i*_ is the *i*^*th*^ layer normalization, 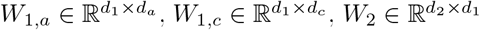 and 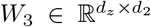 are the weights of the linear layers, while Ψ is the projector’s trainable parameters.

Subsequently, we compute representations for EC numbers by concatenating the sequence and structure representations of proteins under a specific EC number and averaging these concatenated representations:

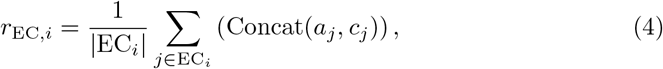

where *r*_EC,*i*_ is the representation of the *i*^*th*^ EC number, EC_*i*_ is the set of enzymes associated with the *i*^*th*^ EC number, |EC_*i*_| is the set’s cardiality, and *a*_*j*_ and *c*_*j*_ are the sequence and structure representations, respectively, of the *j*^*th*^ enzyme under the *i*^*th*^ EC number.

Further, we compute the distance map between the representations of EC numbers utilizing Euclidean distance:

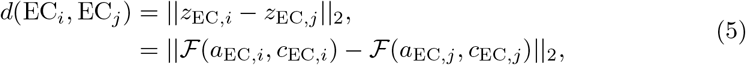

where *d*(EC_*i*_, EC_*j*_) is the Euclidean distance between the projected vectors of the *i*^*th*^ and *j*^*th*^ EC numbers.

During the training phase, we select anchor samples *o* for each EC number and positive samples *p* sharing the same EC number as *o*. For negative samples, we initially choose negative EC numbers with representations that are close to the anchor’s EC number representation in Euclidean distance, then select negative samples from these negative EC numbers. We employ the triplet margin loss [28] for contrastive learning:

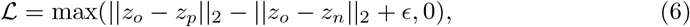

where *ϵ* is the margin for triplet loss (sets as 1 for all experiments), *z*_*o*_ is the projected vector for the anchor sample, *z*_*p*_ is the projected vector for the positive sample, and *z*_*n*_ is the projected vector for the negative sample.

For the EC number prediction phase, we utilize the contrastively trained projector *ℱ* to obtain vectors for enzymes from both the training dataset and test dataset. Subsequently. we compute cluster centers of EC numbers present in the training dataset by averaging the vectors of enzymes associated with each EC number. We then compute the Euclidean distance between the vectors of query enzymes in the test set and vectors of EC numbers. Finally, we predict potential EC numbers utilizing *P* –value algorithm [7] (See Section S1.2).

## Supplementary information

Supplementary Information: Supplementary Materials and Methods and Figs. 1-6 and Tables 1-10.

Supplementary Data 1: Details of 583 annotated enzymes in *Prochlorococcus marinus* MED4 and corresponding prediction outcomes using CLEAN-Contact and CLEAN. Predicted EC numbers were selected using the *P* -value algorithm.

Supplementary Data 2: Details of 1,212 unannotated and unreviewed proteins in *Prochlorococcus marinus* MED4 and corresponding prediction outcomes using CLEAN-Contact. Predicted EC numbers were selected using the *P* -value algorithm.

Supplementary Data 3: Details of enzymes within the New-392 dataset.

Supplementary Data 4: Details of enzymes within the Price-149 dataset.

Supplementary Data 5: Enzymes of New-392 test dataset lacking structures from AlphaFold Protein Structure Database.

Supplementary Data 6: Enzymes of Price-149 test dataset lacking structures from AlphaFold Protein Structure Database.

Supplementary Data 7: Details of EC number prediction outcomes for enzymes within the New-392 dataset using CLEAN-Contact. Predicted EC numbers were selected using the *P* -value algorithm.

Supplementary Data 8: Details of EC number prediction outcomes for enzymes within the New-392 dataset using CLEAN-Contact. Predicted EC numbers were selected using the Max-separation algorithm.

Supplementary Data 9: Details of EC number prediction outcomes for enzymes within the Price-149 dataset using CLEAN-Contact. Predicted EC numbers were selected using the *P* -value algorithm.

Supplementary Data 10: Details of EC number prediction outcomes for enzymes within the Price-149 dataset using CLEAN-Contact. Predicted EC numbers were selected using the Max-separation algorithm.

## Acknowledgments

We extend our gratitude to the Environmental Molecular Sciences Laboratory (EMSL), a DOE Office of Science user facility, for the programmatic funding on project award (10.46936/expl.proj.2022.60535/60008718) that supported the foundational development of the CLEAN-Contact tool. This effort has significantly advanced computational methods for enzyme function prediction and has laid the groundwork for a broad range of scientific applications. This work was also supported by the U. S. Department of Energy, Office of Science, Office of Biological and Environmental Research, under FWP 81832 (NW-BRaVE). PNNL is a multi-program national laboratory operated by Battelle for the DOE under Contract DE-AC05-76RLO 1830. We also appreciate the Predictive Phenomics Initiative (PPI) for providing Laboratory Directed Research and Development (LDRD) funding, which was instrumental in applying the foundational capabilities of CLEAN-Contact to microbial systems. The PPI’s LDRD support facilitated the targeted adaptation of this tool to address the specific challenges within PPI’s research scope, including the enhancement of metabolic modeling and enzyme annotation accuracy. PNNL is a multi-program national laboratory operated by Battelle Memorial Institute for the DOE under Contract DEAC05-76RL01830. This work was also supported by the National Science Foundation (NSF) under Grant #2212465, #2217021, #2217104, #2230111, #2238734, and #2311950 to Qiang Guan.

## Declarations

- Competing interests: The authors declare no competing interests.
- Ethics approval: Not applicable
- Availability of data and materials: All code and data are freely available in open repositories. All codes and data used in training and testing are available at https://github.com/pnnl-predictive-phenomics/clean-contact. CLEAN-Contact is also freely accessible through an easy-to-use web server: https://ersa.guans.cs.kent.edu/.
- Authors’ contributions: J.Z. and Q.G. conceived the study. Y.Y. developed data and codes, and performed all experiments. Y.Y., A.J., S.F., Z.W., J.Z., and Q.G. performed data analyses, and discussed and interpreted all results. Y.Y., Z.W. and C.B. developed the website. Y.Y., M.C., J.Z., and Q.G. wrote the paper.

