## Supplementary Information for "CLEAN-Contact: Contrastive Learning-enabled Enzyme Functional Annotation Prediction with Structural Inference"

Yang et al.

### S1 Supplementary Text

#### S1.1 Data preprocessing

All enzyme amino acid sequences and structures underwent preprocessing prior to training and inference with CLEAN-Contact.

##### S1.1.1 Training dataset

For each protein within the training dataset, proteins lacking structures available in the AlphaFold Protein Structure Database [1] (https://alphafold.ebi.ac.uk/) were excluded from the training dataset. We obtain PDB structures for the rest of the proteins in the training dataset from the AlphaFold Protein Structure Database. The PDB structures are subsequently used to calculate contact maps. The contact map is defined as the distance between $C_\beta - C_\beta$ with a threshold of 8 Å. The contact map was then expanded into three channels with identical values for each channel. We use ResNet-50 [2] with its classification layer removed to acquire structure representation using the expanded contact map. The dimension of the obtained structure representation was $2,048$. Utilizing the ESM-2 model [3], which comprises 36 layers and 3B parameters, we derived sequence representations from the protein's amino acid sequence, resulting in a sequence representation dimension of $2,560$.

##### S1.1.2 Test dataset

In a manner akin to the preprocessing procedures applied to the training dataset, we extracted sequence representations and acquired protein structures from the AlphaFold Protein Structure Database. Contact maps were then derived from these protein structures. However, within the New-392 test dataset, 8 enzymes, and within the Price-149 test dataset, 28 enzymes lacked readily available structures from the AlphaFold Protein Structure Database (Supplementary Data 3 and 4). To address this, we employed all 5 models within AlphaFold2 [4] to generate protein structures for these 36 enzymes. From the 5 generated protein structures for each enzyme, we selected the one with the highest confidence to derive contact maps. Notably, enzymes where an 'X' appeared within the amino acid sequence rendered AlphaFold2 unable to assess structure confidence. In such cases, we utilized the protein structures generated from "model_1". We followed the same procedure mentioned in Section S1.1.1 to obtain the structure and sequence representations for proteins in the test datasets.

#### S1.2 EC number prediction algorithms

There are two EC number selection algorithms that predict EC numbers of query enzymes, the $P$-value EC number selection algorithm and the Max-separation EC number selection algorithm. The $P$-value EC number selection algorithm is based on the statistical significance between the embeddings of enzymes and the embedding of EC numbers. The $P$-value EC number selection algorithm requires the selection of two hyperparameters which define the cutoff for statistical significance to select EC numbers. Higher cutoff values can let the algorithm predict more EC numbers for each query enzyme, potentially discovering previously unknown enzyme functions, while lower cutoff values will result in more accurate predictions. Additionally, with different random seeds, the algorithm could predict slightly different EC numbers for the same query enzyme, especially when the model's confidence of the prediction is extremely low. On the contrary, the Max-separation EC number selection algorithm is less complicated compared the the $P$-value EC number selection algorithm and the Max-separation EC number selection algorithm has no hyperparameter to select. Each time the Max-separation EC number selection algorithm runs, the exact same EC numbers of the same query enzyme will be predicted.

##### S1.2.1 EC number selection using *P*-value

The $P$-value EC number selection algorithm, as proposed by Yu et al. [5], involves several key steps. Initially, a $P$ value was selected, followed by the random sampling of $n$ enzymes from the training dataset, setting a selection threshold $\delta = P \times n$. However, instead of uniformly sampling $n$ enzymes, we assign higher weight to an enzyme whose associated EC numbers contain a smaller number of enzymes:

$$p_e = \frac{1/\max(|\mathrm{EC}_e|)}{\sum_{i \in S} 1/max(|\mathrm{EC}_i|)}, \tag{1}$$

where $p_e$ is the assigned probability of enzyme $e$, $\mathrm{EC}_e$ are all EC number sets including enzyme $e$, $\max(|\mathrm{EC}_e|)$ is the maximum cardinality among all EC number sets involving enzyme $e$, and $S$ is the entire set of enzymes.

Moving forward, we compute the distance map between the randomly sampled $n$ enzymes and EC numbers in the training dataset using vectors encoded by the trained projector $\mathcal{F}$:

$$\begin{aligned} d(\mathrm{EC}_i, e_j) &= ||z_{\mathrm{EC},i} - z_{e,j}||_2, \\ &= ||\mathcal{F}(a_{\mathrm{EC},i}, c_{\mathrm{EC},i}) - \mathcal{F}(a_{e,j}, c_{e,j})||_2, \end{aligned} \tag{2}$$

where $e_j$ is the $j^{th}$ enzyme in the randomly chosen $n$ enzymes, $a$ is the sequence representation, and $c$ is the structure representation. Subsequently, with respect to each EC number, we obtain a set of distances $\mathcal{D}(\mathrm{EC}) = \{d(\mathrm{EC}, e_1), d(\mathrm{EC}, e_2), ..., d(\mathrm{EC}, e_{n-1})\}$, sorted in ascending order.

The next step involves identifying the 10 closest cluster centers of EC numbers to the query enzyme and the corresponding distances between the query enzyme's projected vector to the cluster centers. These 10 EC numbers were denoted as $\mathrm{EC}_k$, with corresponding distances labeled as $d_k$, where $k \in [1, 10]$. $\mathrm{EC}_1$ represents the EC number with the cloesest distance $d_1$ to the query enzyme. We retain the first EC number, which has the smallest distance to the query enzyme, as a prediction result. Subsequently, we iterate over the remaining 9 EC numbers. For each EC number $\mathrm{EC}_k$, we find the index $u$ in $\mathcal{D}(\mathrm{EC}_k)$ where $d_k$ should be inserted such that:

$$d(\mathrm{EC}_k, e_{u-1}) < d_k \leq d(\mathrm{EC}_k, e_{u+1}). \tag{3}$$

We retain $\mathrm{EC}_k$ as a prediction result if the insertion index is smaller than the threshold $\delta$:

$$u \leq \delta. \tag{4}$$

##### S1.2.2 EC number selection using Max-separation

The Max-separation EC number selection algorithm, as introduced by Yu et al. [5], differs from $P$-value algorithm in that it does not need necessitate the selection of hyperparameters $P$ and $n$.

The procedure begins by sorting the distance map between the projected vector of query enzyme and the cluster centers of EC numbers within the training dataset in ascending order. Next, the algorithm selects ten EC numbers $\mathrm{EC}_k$ with the smallest distances $d_k, k \in [1, 10]$ to the query enzyme, computing their average distances:

$$\gamma = \frac{1}{10} \sum_{k=1}^{10} d_k. \tag{5}$$

Following this, the algorithm computes the differences between each distance and the average:

$$q_k = |d_k - \gamma|, k \in [1, 10]. \tag{6}$$

Subsequently, the algorithm computes the differences between adjacent items and computes their average:

$$\begin{aligned} g_k &= |q_k - q_{k-1}|, k \in [2, 10], \\ \bar{g} &= \frac{1}{9} \sum_{i=2}^{10} g_i. \end{aligned} \tag{7}$$

The algorithm proceeds to select the index $i$ for which $g_i > \bar{g}$ and uses $\mathrm{EC}_i$ as the prediction result. In case where no index $i$ satisfies the condition $g_i > \bar{g}$, $\mathrm{EC}_1$ is used as the prediction result.

#### S1.3 Model performance using Max-separation EC number selection algorithm

Besides evaluating CLEAN-Contact using the $P$-value algorithm, we sought to ascertain its comparative performance against CLEAN [5] using the Max-separation EC number selection algorithm. We conducted assessments on both CLEAN-Contact and CLEAN using this algorithm to select potential EC numbers for enzymes in two test datasets. As depicted in Figs. S1 and S2, Precision, Recall, F1-score, and AUC of both CLEAN-Contact and CLEAN were evaluated across the two test datasets. Notably, CLEAN-Contact outperformed CLEAN by exhibiting an average improvement of 10.6% across the two test datasets.

#### S1.4 Model performance when combining representations in different ways

To explore the impact of different combination methods of sequence and structure representations on the performance of CLEAN-Contact, we investigated two additional combination methods beyond the one detailed in the main text (See Section 4). The combination method outlined in the main text, where the two representations are projected to the same dimension and then added together, is denoted as *Addition*.

For the first additional combination method, we initially concatenate the sequence representation and structure representation. This concatenated representation is then passed through the projector:

$$\begin{aligned} z &= \mathcal{F}_\Psi(a, c), \\ &= W_3\left(\mathrm{LN}_2\left(W_2\left(\mathrm{LN}_1(W_1'\mathrm{Concat}(a, c)\right)\right)\right), \end{aligned} \tag{8}$$

where $W_1' \in \mathbb{R}^{d_1 \times (d_a + d_c)}$ is the weight of the first linear layer. This combination method is denoted as $Concat_1$, as the two representations are concatenated before the first linear layer.

For the second additional combination method, we initially pass both representations through the first level linear layers $W_{1,a}$ and $W_{1,c}$ as in the 'Addition' method. Subsequently, we concatenate the outputs of $W_{1,a}$ and $W_{1,c}$ before passing the combined representations through the rest of the neural network:

$$\begin{aligned} z &= \mathcal{F}_\Psi(a, c), \\ &= W_3\left(\mathrm{LN}_2\left(W_2'\left(\mathrm{LN}_1'(\mathrm{Concat}(W_{1,a}a, W_{1,c}c)\right)\right)\right), \end{aligned} \tag{9}$$

where $W_2' \in \mathbb{R}^{d_2 \times (d_1 + d_1)}$ is the weight of the second linear layer. This combination method is denoted as $Concat_2$, reflecting the concatenation of the two representations before the second linear layer.

We proceed to evaluate all three combination methods alongside CLEAN [5] on both the New-392 and Price-149 test datasets. As shown in Tables S1 to S4, it is evident that, regardless of the approach used to combine sequence and structure representations, they consistently outperformed CLEAN, which lacks structural information.

Notably, among the three combination methods, the *Addition* consistently demonstrated superior performance. We attribute this outcome to the principles of residual learning, known to facilitate optimization processes [2].

#### S1.5 Analysis of model uncertainty

We harness Gaussian Mixture Model (GMM) [6] to quantify the model's confidence during prediction. Initially, we randomly sampled 500 different EC numbers. Subsequently, we compute the distances between the cluster centers of these EC numbers and the combined representations of the associated enzymes. The distribution of enzyme counts versus distances forms a "same-EC" Gaussian distribution, representing the enzymes with correct EC numbers (Left peak in Fig. S3). Following Section 4.2, we sampled negative enzymes for these 500 EC numbers and computed distances between the cluster centers of these EC numbers and those of the sampled negative enzymes. The distribution of negative enzyme counts versus distances form a "different-EC" Gaussian distribution, representing the enzymes with incorrect EC numbers (Right peak in Fig. S3). Finally, we employ a 2-component GMM to fit these two Gaussian distributions, with one component representing the "same-EC" Gaussian distribution and the other representing the "different-EC" Gaussian distribution.

We begin by computing the distance between the combined representation of a test enzyme and that of the predicted EC number. Subsequently, we calculate the density of the 2-component fitted GMM for this distance. Utilizing the density of the component corresponding to the "same-EC" distribution from the fitted GMM as the prediction confidence, we evaluate the confidence level for this prediction result. Following this, we assess the performance of CLEAN-Contact across increasing cumulative confidence levels on the merged test dataset. As shown in Figs. S4 and S5, the Precision and Recall of CLEAN-Contact show improvement with higher cumulative confidence, consistently outperforming CLEAN across all cumulative confidence levels.

Additionally, we employ the fitted GMM to address overprediction concerns. Specifically, we restrict predictions to the 4th-level EC number only when the prediction confidence exceeds 0.5; otherwise, we predict the 3rd-level EC number. As illustrated in Fig. S6, this adaptive prediction strategy mitigates overprediction issues by favoring the prediction of 3rd-level EC numbers at lower confidence levels. Remarkably, when employing this adaptive prediction strategy, CLEAN-Contact yields more true positives compared to exclusively predicting 4th-level EC numbers across all scenarios. Additionally, CLEAN-Contact consistently outperforms CLEAN in terms of predictive accuracy.

#### S1.6 Selection of pretrained model for structure representation extraction

To determine the optimal model for extracting structure representations, we trained and evaluated four CLEAN-Contact variants employing different CNNs: three ResNet-based models [2] (ResNet-18, ResNet-50, and ResNet-101) and one vision transformer-based model (SwinV2-B [7]). While ResNet-18 has the lowest computational cost, it also demonstrates the lowest performance in ImageNet classification tasks [8]

(Table S5). ResNet-50, ResNet-101, and SwinV2-B exhibit comparable performance in these tasks, while the number of parameters ranges from 25.6M (ResNet-50) to 87.9M (SwinV2-B) (Table S5). Our experiments on the merged test dataset revealed that ResNet-based models in CLEAN-Contact showed similar performance, with ResNet-50 slightly outperforming others (Table S6). Interestingly, SwinV2-B variant of CLEAN-Contact underperformed compared to the ResNet-based models (Table S6). These findings suggested that ResNet-50 offers the optimal balance, maximizing the extraction of useful information from protein contact maps while maintaining computational efficiency.

#### S1.7 Selection of optimal number of samples for contrastive learning

To study the impact of sample quantities for contrastive learning on the performance of CLEAN-Contact, we conducted a series of experiments with different numbers of anchor, positive, and negative samples. We trained models with 1, 2, and 4 samples for each sample type and adhered to the constraints imposed by the triplet margin loss function. The triplet margin loss function requires an equal number of positive and negative samples. Additionally, it requires that the number of positive samples must be either one, equal to the number of anchor samples, or greater than the number of anchor samples. We observed an inverse relationship between sample quantity and model performance. Notably, increasing the number of positive and negative samples led to a more significant decrease in performance compared to increasing the number of anchor samples alone (Table S7).

#### S1.8 Analysis of computational cost

Computational cost is another important factor in evaluating the performance of the classification model. We evaluated the computational cost of CLEAN-Contact and CLEAN, focusing on their inference steps. For CLEAN-Contact, this includes generating sequence representations using ESM-2, structure representations using ResNet-50, and predicting EC numbers of query enzymes using the contrastively learned model. CLEAN's inference steps include obtaining representations through ESM-1b and making predictions. We quantified computational complexity using Giga Floating Operations (GFLOPs) and time consumption for each step.

To ensure a fair comparison, we used the average protein sequence length from our test dataset (439 amino acids) as a benchmark, since the computational cost of ESM-1b, ESM-2, and ResNet-50 are input-size dependent. We excluded the cost of generating PDB structures for proteins lacking entries in the AlphaFold Protein Structure Database [1].

Our analysis revealed that CLEAN-Contact requires only an additional 0.1776 seconds per enzyme compared to CLEAN (Table S8) while achieving significant performance improvement over CLEAN, scoring more than 20% improvement in several benchmark metrics.

##### S1.9 Cross-validation evaluation

To rigorously evaluate model performance and address potential overfitting concerns, we conducted a 5-fold cross-validation for both CLEAN-Contact and CLEAN. We then assessed each of these models on our test datasets (New-392 and Price-149), calculating the mean and standard deviations of performance metrics across the five runs. Notably, even when trained only on 80% of the training dataset, CLEAN-Contact consistently outperformed CLEAN across both test datasets (Table S9 and Table S10).

#### S2 Supplementary Figures

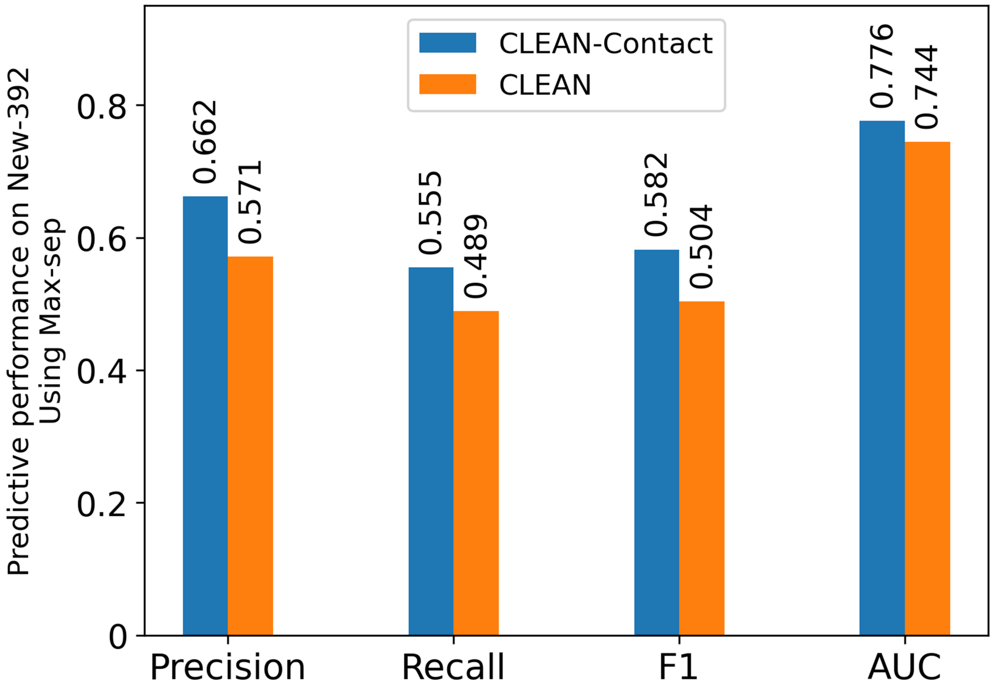


**Fig. S1** Evaluation of predictive performance between CLEAN-Contact and CLEAN on the New-392 test dataset, evaluated using Precision, Recall, F1-score, and AUC metrics. The Max-separation EC number selection algorithm was employed for selecting potential EC numbers.

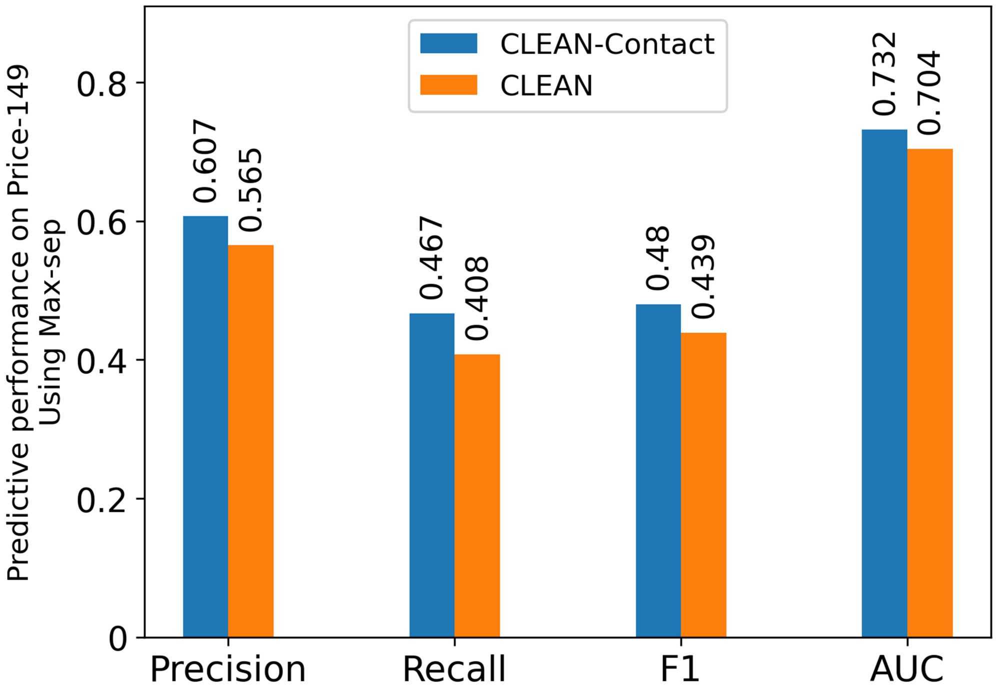


**Fig. S2** Evaluation of predictive performance between CLEAN-Contact and CLEAN on the Price-149 test dataset, evaluated using Precision, Recall, F1-score, and AUC metrics. The Max-separation EC number selection algorithm was employed for selecting potential EC numbers.

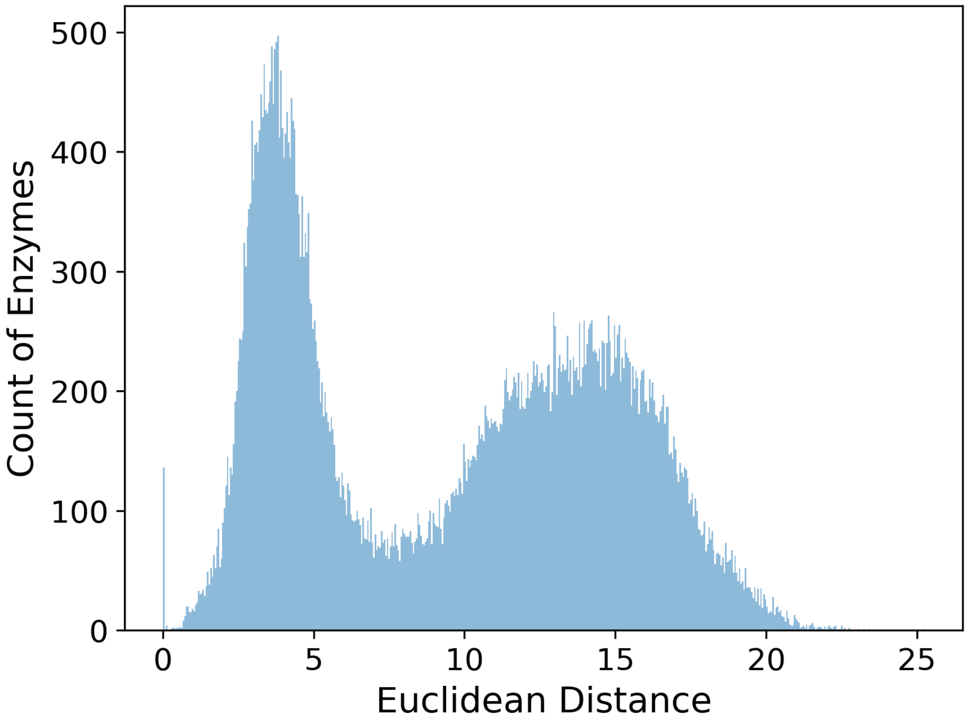


**Fig. S3** Histogram showing the distribution of randomly sampled enzymes plotted against the Euclidean distance between the enzyme's combined representation and the cluster centers of EC numbers. The left peak illustrates the distribution when sampled enzymes correspond to their respective EC numbers while the right peak illustrates the distribution when sampled enzymes do not match their corresponding EC numbers.

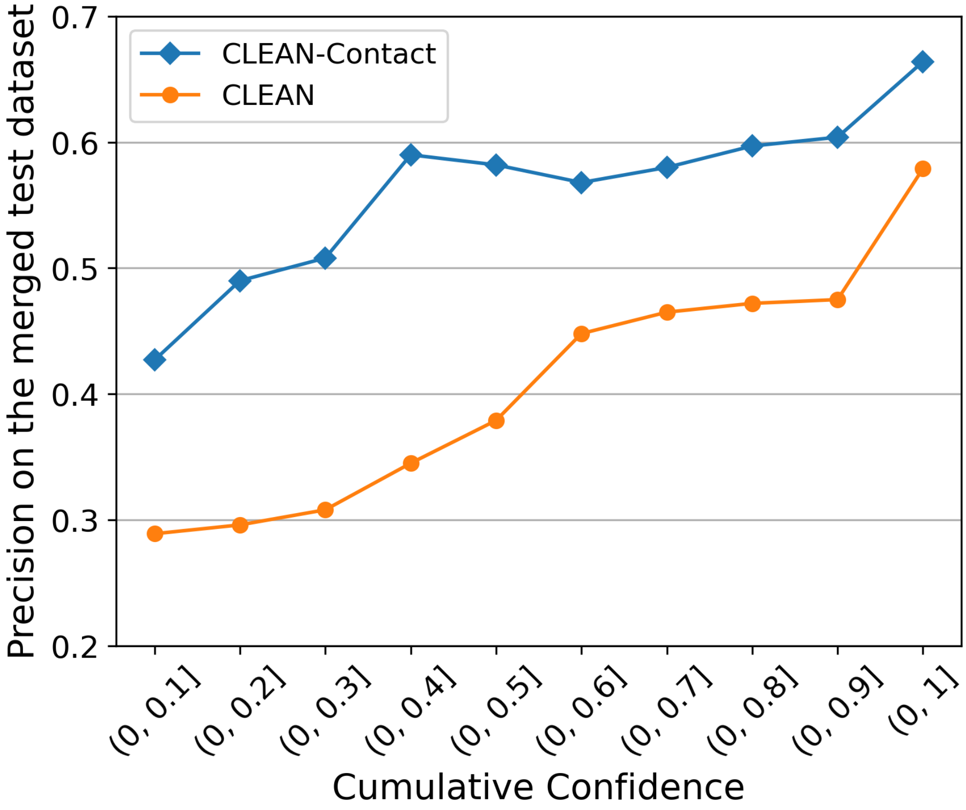


**Fig. S4** Evaluation of predictive performance comparing CLEAN-Contact and CLEAN on the merged test dataset, evaluated using the Precision metric. This evaluation correlates with the ascending cumulative confidence derived from the fitted Gaussian Mixture Model.

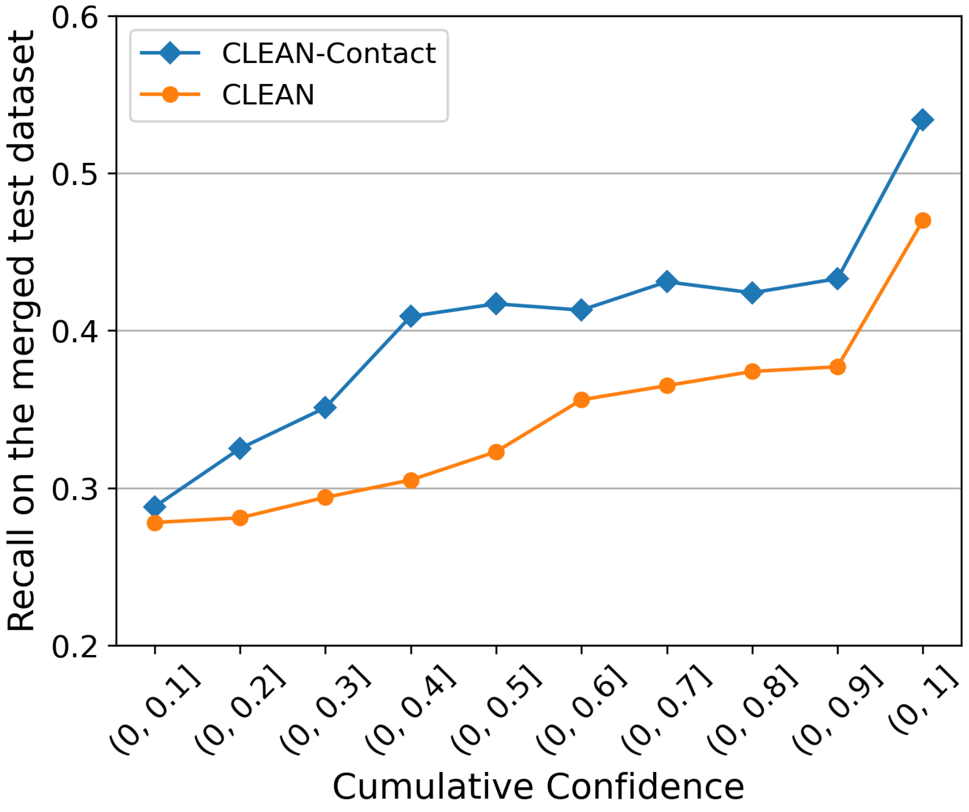


**Fig. S5** Evaluation of predictive performance comparing CLEAN-Contact and CLEAN on the merged test dataset, evaluated using the Recall metric. This evaluation correlates with the ascending cumulative confidence derived from the fitted Gaussian Mixture Model.

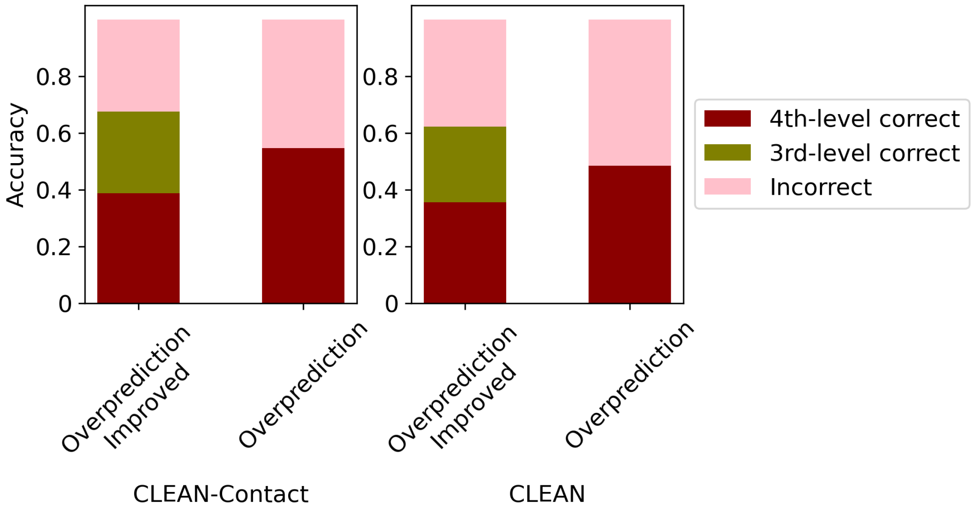


**Fig. S6** Mitigating overprediction issue through an adaptive prediction strategy, evaluated via Accuracy. Left: predictive performance of CLEAN-Contact is depicted when employing two strategies: predicting 4th-level EC numbers solely at high prediction confidence and predicting 3rd-level EC numbers otherwise (left bar: "Overprediction Improved"), versus predicting 4th-level EC numbers at all times (right bar: "Overprediction"). Right: predictive performance of CLEAN under the same two strategies: predicting 4th-level EC numbers solely at high confidence and predicting 3rd-level EC numbers otherwise (left bar: "Overprediction Improved"), versus predicting 4th-level EC numbers at all times (right bar: "Overprediction").

#### S3 Supplementary Tables

| Combination | Precision | Recall | F1 | AUC |
|---|---|---|---|---|
| *Addition* | 0.652 | 0.555 | 0.566 | 0.777 |
| $Concat_1$ | 0.616 | 0.525 | 0.528 | 0.761 |
| $Concat_2$ | 0.613 | 0.549 | 0.557 | 0.774 |
| CLEAN | 0.561 | 0.509 | 0.504 | 0.753 |

**Table S1** Evaluation of predictive performance among various combinations of sequence and structure representations alongside CLEAN on the New-392 test dataset. Predicted EC numbers were selected using the $P$-value EC number selection algorithm.

| Combination | Precision | Recall | F1 | AUC |
|---|---|---|---|---|
| *Addition* | 0.662 | 0.555 | 0.582 | 0.776 |
| $Concat_1$ | 0.636 | 0.507 | 0.528 | 0.753 |
| $Concat_2$ | 0.626 | 0.497 | 0.524 | 0.748 |
| CLEAN | 0.571 | 0.489 | 0.504 | 0.744 |

**Table S2** Evaluation of predictive performance among various combinations of sequence and structure representations and CLEAN on the New-392 test dataset. Predicted EC numbers were selected using the Max-separation EC number selection algorithm.

| Combination | Precision | Recall | F1 | AUC |
|---|---|---|---|---|
| *Addition* | 0.621 | 0.513 | 0.525 | 0.756 |
| $Concat_1$ | 0.562 | 0.480 | 0.484 | 0.739 |
| $Concat_2$ | 0.563 | 0.441 | 0.453 | 0.720 |
| CLEAN | 0.531 | 0.434 | 0.452 | 0.717 |

**Table S3** Evaluation of predictive performance among various combinations of sequence and structure representations and CLEAN on the Price-149 test dataset. Predicted EC numbers were selected using the $P$-value EC number selection algorithm.

| Combination | Precision | Recall | F1 | AUC |
| --- | --- | --- | --- | --- |
| *Addition* | 0.607 | 0.467 | 0.480 | 0.732 |
| $Concat_1$ | 0.560 | 0.441 | 0.447 | 0.718 |
| $Concat_2$ | 0.507 | 0.434 | 0.432 | 0.716 |
| CLEAN | 0.565 | 0.408 | 0.439 | 0.704 |

**Table S4** Evaluation of predictive performance among various combinations of sequence and structure representations and CLEAN on the Price-149 test dataset. Predicted EC numbers were selected using the Max-separation EC number selection algorithm.

| | Accuracy (Top-1) | Parameter # |
| --- | --- | --- |
| ResNet-18 | 69.758% | 11.7M |
| ResNet-50 | 80.858% | 25.6M |
| ResNet-101 | 81.886% | 44.5M |
| SwinV2-B | 84.112% | 87.9M |

**Table S5** Comparison of benchmark performance on computer vision classification task and computational cost between commonly used convolutional neural networks (CNNs).

| Model | Precision | Recall | F1-score | AUC |
| --- | --- | --- | --- | --- |
| ResNet-50 & ESM-2 | 0.634 | 0.562 | 0.563 | 0.780 |
| ResNet-18 & ESM-2 | 0.593 | 0.515 | 0.516 | 0.757 |
| ResNet-101 & ESM-2 | 0.639 | 0.566 | 0.560 | 0.778 |
| SwinV2-B & ESM-2 | 0.370 | 0.246 | 0.269 | 0.622 |

**Table S6** Performance evaluation of CLEAN-Contact variants on the merged dataset. Different pretrained computer vision models (ResNet-50, ResNet-18, ResNet-101, and SwinV2-B) were used to extract structure representations from protein contact maps.

| # of Anchor Samples | # of Positive (Negative) Samples | Precision | Recall | F1-score | AUC |
|---|---|---|---|---|---|
| 1 | 1 | 0.634 | 0.562 | 0.563 | 0.78 |
| 1 | 2 | 0.626 | 0.542 | 0.552 | 0.77 |
| 1 | 4 | 0.601 | 0.513 | 0.521 | 0.757 |
| 2 | 1 | 0.632 | 0.51 | 0.515 | 0.754 |
| 2 | 2 | 0.631 | 0.527 | 0.542 | 0.763 |
| 4 | 1 | 0.632 | 0.53 | 0.536 | 0.764 |
| 4 | 4 | 0.62 | 0.51 | 0.528 | 0.754 |

**Table S7** Evaluation of performance of CLEAN-Contact on the merged test dataset with different number of anchor, positive, and negative samples. The performance was measured using Precision, Recall, F1-score, and AUC.

| | CLEAN-Contact | | CLEAN | |
|---|---|---|---|---|
| | GFLOPs | Inference time (Nvidia A5000) | GFLOPs | Inference time (Nvidia A5000) |
| ESM-2 | 46,035 | 0.2379 sec | N/A | N/A |
| ESM-1b | N/A | N/A | 10,846 | 0.0682 sec |
| ResNet-50 | 16.164 | 0.0078 sec | N/A | N/A |
| Prediction | 0.003 | 0.0009 sec | 0.001 | 0.0008 sec |
| Total | 46,051.167 | 0.2466 sec | 10,846.001 | 0.0690 sec |

**Table S8** Comparison of computational cost for inference of CLEAN-Contact and CLEAN on one protein using Giga Floating Operations (GFLOPs) and inference time on Nvidia A5000 GPU. The computational costs of the pretrained models, which are ESM-2, ESM-1b, and ResNet-50, are calculated using the average length of amino acid sequences in the test datasets, which is 439 amino acids.

| Model | Precision | Recall | F1-score | AUC |
|---|---|---|---|---|
| CLEAN-Contact | $0.642 \pm 0.014$ | $0.544 \pm 0.01$ | $0.55 \pm 0.005$ | $0.771 \pm 0.005$ |
| CLEAN | | | | |

**Table S9** Comparison of 5-fold cross-validation result between CLEAN-Contact and CLEAN on the New-392 test dataset. Results show mean values with standard deviations.

| Model | Precision | Recall | F1-score | AUC |
|---|---|---|---|---|
| CLEAN-Contact | $0.592 \pm 0.006$ | $0.487 \pm 0.006$ | $0.496 \pm 0.003$ | $0.743 \pm 0.003$ |
| CLEAN | | | | |

**Table S10** Comparison of 5-fold cross-validation result between CLEAN-Contact and CLEAN on the Price-149 test dataset. Results show mean values with standard deviations.
